# Connectome preprocessing by Consensus Clustering increases separability in group neuroimaging studies

**DOI:** 10.1101/348110

**Authors:** Javier Rasero, Jesus M Cortes, Daniele Marinazzo, Sebastiano Stramaglia

## Abstract

One of the biggest challenges in preprocessing pipelines for neuroimaging data is to increase the signal-to-noise ratio of the data which will be used for subsequent analyses. In the same line, we suggest in the present work that the application of consensus clustering for brain connectivity matrices to find subgroups of **subjects** can be a valid additional”connectome processing” step helpful to reduce intra-group variability and therefore increase the separability of distinct classes. In addition, by partitioning the data before any group comparison, we demonstrate that unique regions within each cluster arise and bring new information that could be relevant from a clinical point of view.

## I. INTRODUCTION

In the supervised classification of human connectome data [1], subjects are usually grouped based on high-level clinical categories (e.g., patients and controls), and typical approaches aim at deducing a decision function from the labeled training data, see e.g.[2]. Likewise, unsupervised analysis are also usually performed such that one is blind to any phenotypic factors and is more interested in finding subgroups of subjects/features with similar characteristics. There exist in the literature a vast number of clustering algorithms dealing with this issues (see for example [3] and references in). The emergence of substructures underlying the data is due to the fact that in general the population of healthy subjects (as well as those of patients) is usually highly heterogeneous. Consequently, stratification of groups may be a useful preprocessing stage, so that the subsequent supervised analysis might exploit the knowledge of the structure of data. A convenient strategy for stratification of groups involves using phenotypic variables of subjects when available. More interestingly, stratification may rely on the measured variables, like the human connectome data itself, by application of clustering algorithms that find natural groupings in the data.

An effective supervised approach, named Multivariate Distance Matrix Regression (MDMR), has been proposed in [4] for the analysis of gene expression patterns; it tests the relationship between variation in a distance matrix and predictor information collected on the samples whose gene expression levels have been used to construct the matrix. The same method was also applied to the cross-group analysis of brain connectivity matrices [5], as an alternative to the common method used in connectome-wide association studies, i.e. mass-univariate statistical analyses, in which the association with a phenotypic variable of each entry of the brain connectivity matrix, across subjects, is tested. Whilst MDMR has found wide application, see e.g. [6], its findings may be certainly affected by the heterogeneity of classes.

Recently an unsupervised method [7], rooted on the notion of *consensus* clustering [8], has been developed for community detection in complex networks [9], when a connectivity matrix is associated to each item to be classified; in this method, the different features, extracted from connectivity matrices, are not combined in a single vector to feed the clustering algorithm; rather, the information coming from the various features are combined by constructing a *consensus* network [8]. Consensus clustering is commonly used to generate stable results out of a set of partitions delivered by different clustering algorithms (and/or parameters) applied to the same data [10]; in [7], instead, the consensus strategy was used to combine the information about the data structure arising from different features so as to summarize them in a single consensus matrix, which not only provides a partition of subjects in communities, but also a geometrical representation of the set of subjects. It has been shown that it is an effective technique for disentangling the heterogeneity that is inherent to many conditions, and to the cohort of controls. The main similarity with MDMR is that in both methods a distance matrix in the space of subjects is introduced.

The purpose of this work is to propose the consensus clustering approach in [7] as a preprocessing stage of MDMR in exploratory analysis, so as to cope with the heterogeneity of subjects. First, we will test the robustness of our consensus clustering with respect to the introduction of heterogeneity in the data provided by means of different affects of noise. Second, we will show that extracting the natural classes present in data and subsequently performing the supervised analysis between the subgroups found by consensus clustering, allows to identify variables whose pattern is altered in group comparisons, which are not identified when the groups are used as a whole. As a result, the proposed approach leads to an increase in separability.

## II. MATERIALS AND METHODS

### A. Materials

The robustness of the consensus clustering algorithm to disentangle the real structure of data was tested using simulated data for two groups of 15 subjects each, generated using *simTB* [11], a toolbox written in MATLAB that allows to simulate functional magnetic resonance imaging (fMRI) datasets under a model of spatiotemporal separability. Given the number of sources/components (*n*_*C*_), repetition time (TR) and a hemodynamic model, the program yields the time course profile subject to the experimental design (block- and/or event-related) provided. Our scenario consists of *n*_*C*_ = 20 components and *TR* = 2s, such that in a event-related experiment the first 10 components of the cohort of group 1 have high probability (90%) of becoming activated, whereas for the remaining components the activation is rare (10 %). For subjects of group 2 the situation is the opposite. This scenario could be then considered as the same group of subjects performing two orthogonal task, where different components get involved.

**Table I:** Demographic information for LA5C dataset

|  | Sex (M/F) | Age | Framewise displacement (FWD) |
| --- | --- | --- | --- |
| HC (N=118) | 64/54 | $31.56 \pm 8.83$ | $0.17 \pm 0.15$ |
| ADHD (N=40) | 21/19 | $32.05 \pm 10.4$ | $0.15 \pm 0.11$ |
| BP (N=47) | 26/21 | $35.47 \pm 9.16$ | $0.19 \pm 0.13$ |
| SCH (N=50) | 38/12 | $36.46 \pm 8.88$ | $0.25 \pm 0.20$ |

On the other hand the advantages of using the consensus clustering method as a preprocessing step were explored on two real and public neuroimaging datasets. The first dataset comprises Healthy Controls (HC) subjects and several pathologies: Attention-deficit hyperactivity disorder (ADHD), Bipolar Disorder (BD), Schizophrenia (SCH). Such a dataset can be found in the UCLA Consortium for Neuropsychiatric Phenomics LA5C Study from the OpenfMRI database with accession number ds000030 [12], from which we used resting functional MRI (rfMRI) of 255 subjects: 40 ADHD, 47 BD, 50 SCH and 118 HC. Demographics information about this cohort can be found in table I. Data were preprocessed with FSL (FMRIB Software Library v5.0). All volume images were corrected for motion, after which slice timing correction was applied to correct for temporal alignment. All voxels were spatially smoothed with a 6mm FWHM isotropic Gaussian kernel and a band pass filter was applied between 0.01 and 0.08 Hz after intensity normalization. In addition, linear and quadratic trends were removed. We next regressed out the motion time courses, the average CSF signal and the average white matter signal. Global signal regression was not performed. Data were transformed to the MNI152 template, such that a given voxel had a volume of 3mm × 3 mm × 3mm. Finally we obtained 278 time series, each corresponding to an anatomical region of interest (ROI), by averaging the voxel signals according to the functional atlas described in [13]. Finally, a 278 × 278 matrix of Pearson coefficients amongst time series for each subject was obtained.

As for the second dataset, it corresponds to unprocessed Resting-State fMRI signal for a population of healthy controls and Autism Spectrum Disorder (ASD) subjects that can be found in the Autism Brain Imaging Data Exchange (ABIDE) repository [14], an initiative that aims at collecting data of this disorder from laboratories around the world to accelerate the understanding of its neural bases. Since it is unclear the effect that different scanner machines can have on the observable results, we decided to avoid this issue by considering only a sample of 75 subjects for each group acquired at the same site (NYU). For this subset, we matched age and sex between samples (Wilcoxon rank sum *p* = 0.27 and χ^2^ test *p* = 0.29 respectively). Preprocessing of the data was done using FSL, AFNI and Matlab. Firstly, slice-time correction was applied. Then each volume was aligned to the middle volume to correct for head motion artefacts followed by intensity normalization. We next regressed out 24 motion parameters, the average cerebrospinal fluid (CSF) and the average white matter signal. A band pass filter was applied between 0.01 and 0.08 Hz and linear and quadratic trends were removed. All voxels were spatially smoothed with a 6 mm full width at half maximum (FWHM). Finally, FreeSurfer software was used for brain segmentation and cortical parcellation. A 86 region atlas was generated with 68 cortical regions from Desikan-Killiany Atlas (34 in each hemisphere) and 18 subcortical regions (9 in each hemisphere: Thalamus, Caudate, Putamen, Pallidum, Hippocampus, Amygdala, Accumbens, VentralDC and Cerebellum). Each subject parcellation was projected to individual functional data and the mean functional time series of each region was computed.

### B. Consensus clustering method

Given a set of matrices of distance among subjects, the consensus clustering proposed in [7] can be summarised as follows: (i) clusterise each distance matrix using a known clustering algorithm, (ii) build the consensus network from the corresponding partitions and (iii) extract groups of subjects by finding the communities of the consensus network thus obtained.

Regarding the distance matrices calculation in the case of fMRI data, let us consider *m* subjects such that each one has a *N* × *N* matrix of features, that can for example represent the brain functional connectivity matrix. We will denote this matrix as {**A**(**i**,**j**)_α_}, where α = 1,…,*m* and *i*, *j* = 1,…, *N*. For each row *i*, we build a distance matrix for the set of subjects as follows. Consider a pair of subjects α and β and consider the corresponding patterns {**A**(**i**, :)α} and {**A**(**i**, :)β} let r be their Pearson correlation. As the distance between the two subjects, for the node i, we take *d*_*αβ*_ = 1 – *r*; other choices for the distance can be used, like, e.g., 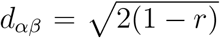 where *r* is the Pearson correlation. The m × m distance matrix *d*_*αβ*_ corresponding to row *i* will be denoted by **D**_i_, with *i* = 1,…, *N*. The set of **D** matrices may be seen as corresponding to layers of a multilayer network [15], each brain node providing a layer.

Each distance matrix **D**_i_ is then partitioned into *k* groups of subjects using k-medoids method [16]. Subsequently, an *m* × *m* consensus matrix **C** is evaluated: its entry *C*_*αβ*_ indicates the number of partitions in which subjects α and β are assigned to the same group, divided by the number of partitions N. Eventually the consensus matrix is averaged over *k* ranging in the interval (2-20) so as to fuse, in the final consensus matrix, information about structures at different resolutions, see [7].

The consensus matrix, obtained as explained before, is eventually partitioned in communities by modularity maximization, with the consensus matrix **C** being compared against the ensemble of all consensus matrices one may obtain randomly and independently permuting the cluster labels obtained after applying the k-medoids algorithm to each of the set of distance matrices. More precisely, a modularity matrix is evaluated as

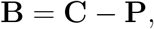

where **P** is the expected co-assignment matrix, uniform as a consequence of the null ensemble here chosen, obtained repeating many times the permutation of labels; the modularity matrix **B** is eventually submitted to a modularity optimization algorithm to obtain the output partition by the proposed approach (we used the Community Louvain routine in the Brain Connectivity Toolbox [17], which admits modularity matrices instead of connectivity matrices as input).

### C. Multivariate Distance Matrix Regression

The cross-group analysis of brain connectivity matrices has been performed using the Multivariate Distance Matrix Regression (MDMR) approach as described in [4, 5].

For a fixed brain node i, the distance between connectivity patterns of i with the rest of the brain was calculated per pair of subjects (u,v) -by calculating Pearson correlation between connectivity vectors of subject pairs-, thus leading to a distance matrix in the subject space for each i investigated. In particular, the following formula was applied

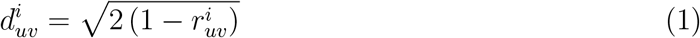

where 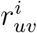 is the Pearson correlation between connectivity patterns of i for subjects u and v. Next, MDMR was applied to perform cross-group analysis as implemented in R [18].

MDMR yielded a pseudo-F estimator (analogous to that F-estimator in standard ANOVA analysis), which addresses significance due to between-group variation as compared to within-group variations [19]. In the particular case when the only regressor variable is categorical (i.e. the group label), given a distance matrix, one can calculate the total sum of squares as

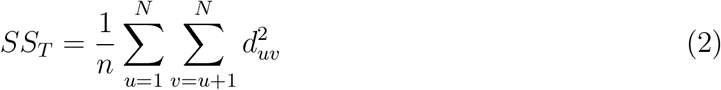

with N being the total number of subjects. Notice that, from here on, we will consider Thus, we got a different for each module i. Similarly, the within-group sum of squares can be written as

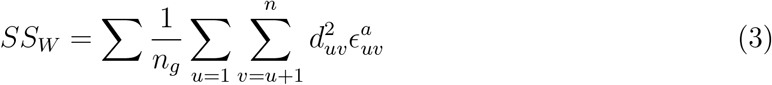

where is the number of subjects per group and a variable equal to 1 if subjects u and v belong to group g and 0 otherwise. The between-group variation is simply, which leads to a pseudo-F statistic that reads

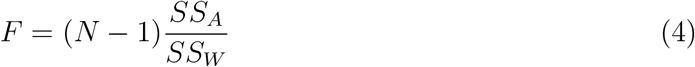

where m is the number of groups. As it was acknowledged in [4], the pseudo-F statistic is not distributed like the usual Fisher’s F-distribution under the null hypothesis. Accordingly, we randomly shuffled the subject indexes and computed the pseudo-F statistic for each time. A p-value is computed by counting those pseudo F-statistic values from permuted data greater than that from the original data respect to the total number of performed permutations. Nevertheless, in our cross-group analyses, in addition to group label, age, sex and frame-wise displacement (FWR) were also considered as covariates since their possible confounding effect in distance variation between subjects. Finally, we controlled for I errors by False discovery rate corrections and set significance threshold at 1%.

### D. The proposed approach

The application of MDMR, to identify altered patterns, is hampered by the heterogeneity of subjects in the same group (healthy or patients). Therefore we propose here the use of the consensus clustering approach as a preprocessing stage, to extract the natural classes present in the group of controls (and/or in the group of patients); subsequently the supervised analysis of MDMR is to be performed between the pairs of subgroups found by consensus clustering, so as to identify variables whose pattern is altered in subgroups comparisons, whilst they are not identified as significantly altered when the groups are used as a whole.

## III RESULTS

### A. Simulated data

Our approach is based on the benefit of gain cluster information from each node instead of using the whole brain. We can visualise this in figure 1, where the group reconstruction provided by our method is compared with the one yielded by considering the whole correlation matrix as pattern connectivity from which calculate the distance matrix, so the average is only carried out over different resolutions *κ* when applying k-medoids.

We also assessed the robustness of our method in group reconstruction when gaussian noise is added to the time series of group 1 (*TS*_1_) and group 2 (*TS*_2_), that can be written as follows

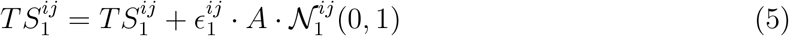

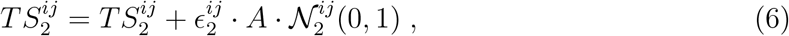

for *i* =1… 15 subjects and *j* = 1… 20 components. *A* is the amplitude of the perturbations and ∊_ij_ is a binary matrix allowing us to play with different noise configuration scenarios.

In a first scenario, different values of the noise amplitude A={0.1,0.3,0.5} were applied to all components and subjects 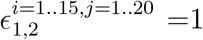. As we can see in figure 2, only for large amplitude, noise starts to dominate so much that the consensus algorithm renders group more compact and mix them together making it incapable of distinguishing between groups perfectly

**Fig. 1:**
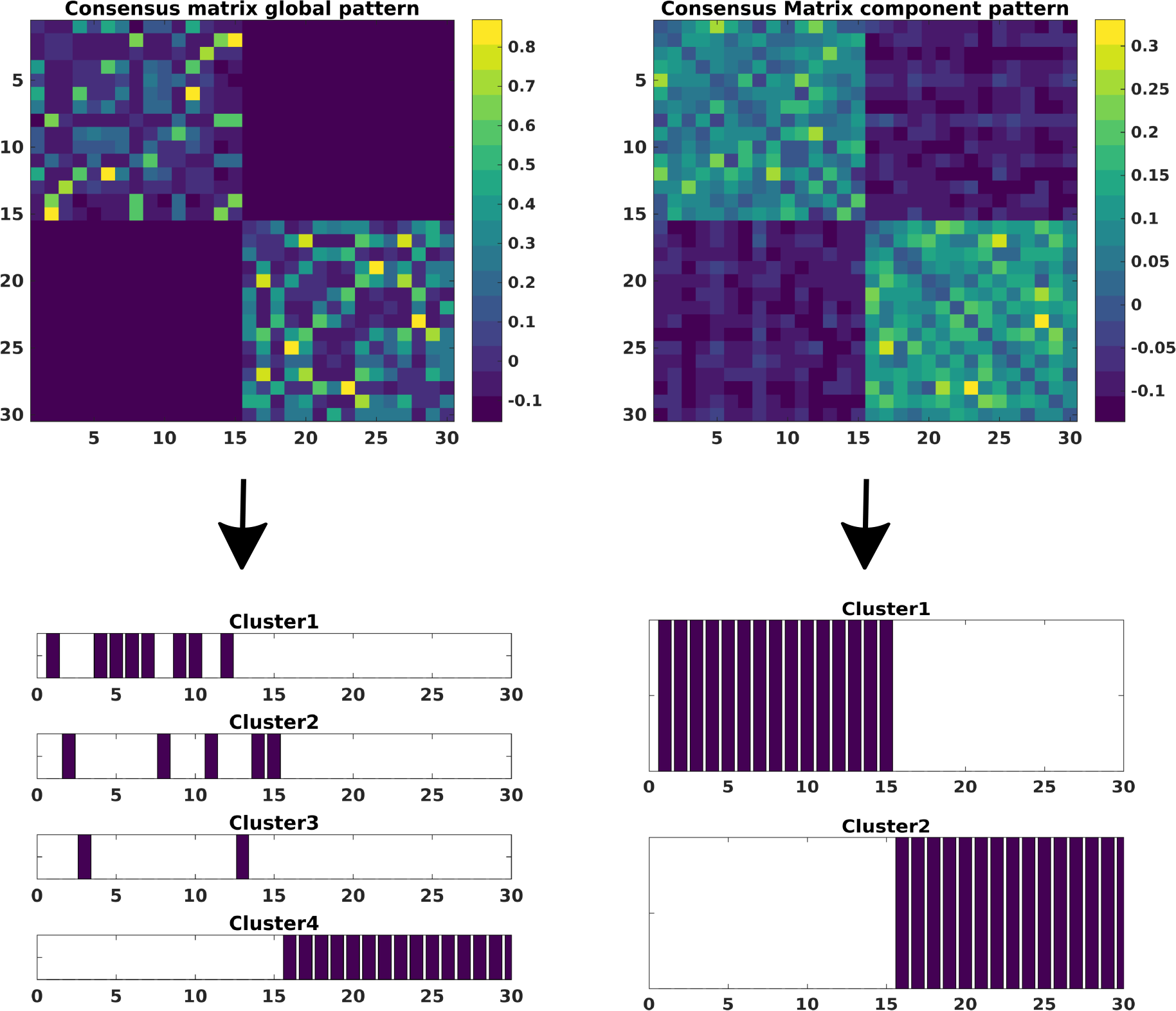
Comparison of group reconstruction obtained after the application of Louvain community detection to the modularity matrix *B* provided by our method using the whole pattern connectivity matrix (upper left) or the node pattern connectivity (upper right)

In a second scenario, variation of noise according to subsets of components perturbed was studied. In order to amplify this effects, we restrict noise amplitudes to a fixed large value A = 0.5. Four subsets of components affected by noise were taken *j* = {(1), (1… 3), (1… 6), (1… 10)}. In addition, in order to add more complexity, we considered the application of noise to 5 subjects of group 1 and 10 of group 2 respectively. As a consequence, this effect is more notorious in group 2, since for the first 10 components the activity is lower and therefore more sensitive to noise. When applied to a different number of components, this leads to a decrease of intra-group distance of subjects affected (yellowish areas in the consensus matrices of figure 3) and also a slight approximation between groups as a consequence of the synchronisation amongst the noised components. Nevertheless, the consensus clustering is pretty robust when reconstructing both groups

**FIG. 2:**
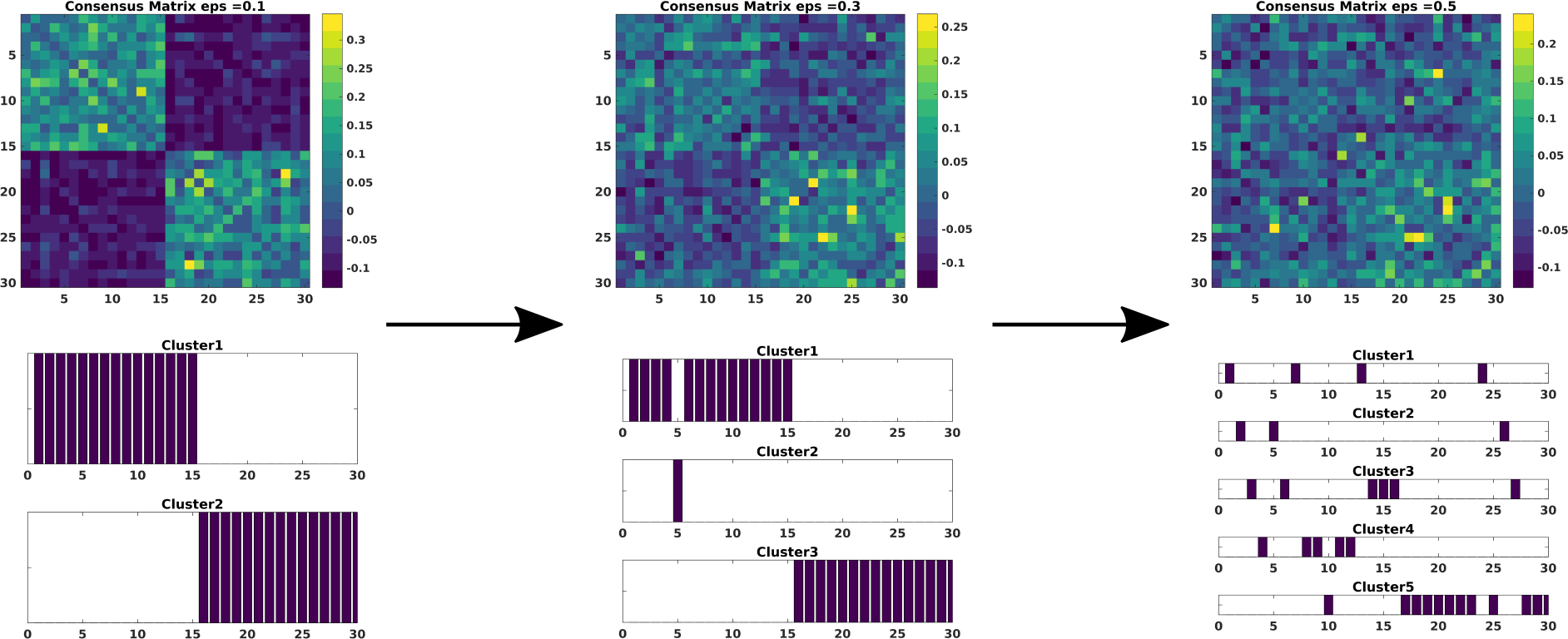
Comparison of the evolution of the modularity matrix B and group reconstruction when changing the amplitude noise that affects to all the simulated time series for both groups

**FIG. 3:**
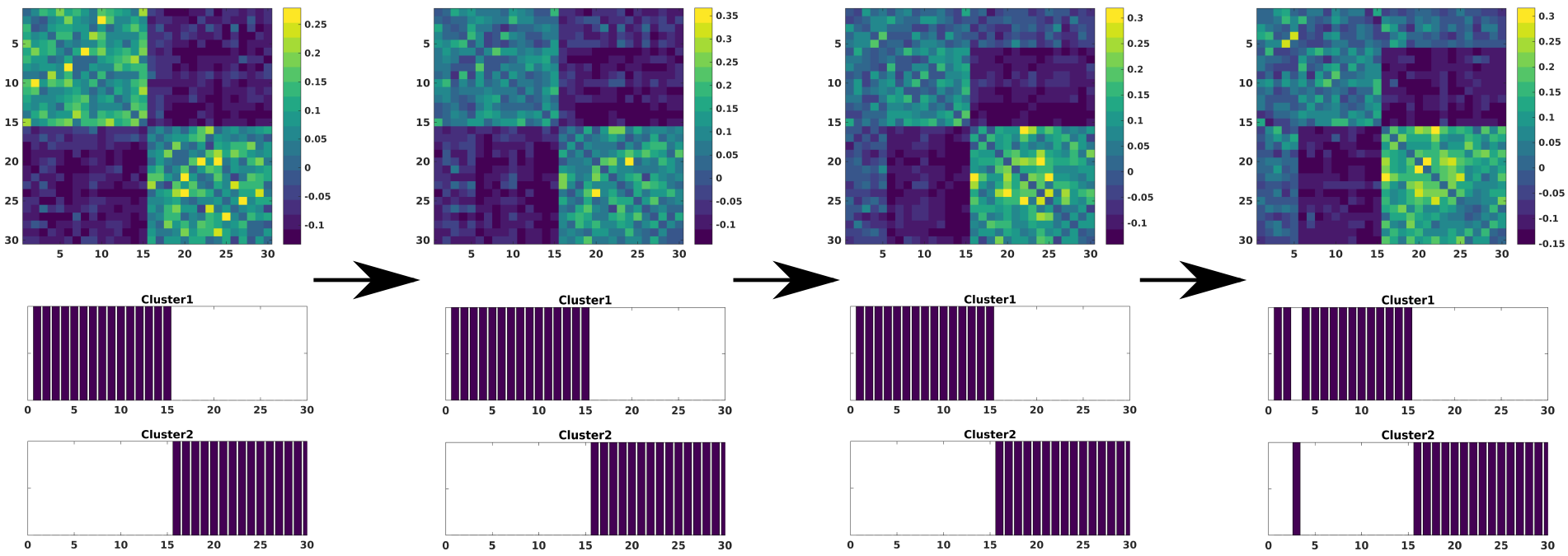
Comparison of the evolution of the modularity matrix *B* and group reconstruction when changing the number of components affected by large amplitude noise

Finally, variation of number of subjects was taken into account. In order to simplify this scenario, we considered that noise only affected group 2 and the first 10 components with a large amplitude *A* = 0.5. The subset of number of subjects are *i* = {(1), (1… 5), (1… 10), (1… 15)}. The consensus turns out to be again robust when reconstructing both groups. For this case, the effect of changing the number of subjects of group 2 affected by noise makes this group more compact as more subjects are involved since noise makes them look more similar. This also makes these subjects of group 2 be more similar with those of group 1 for the first 10 components, but not enough to mix them together to render them indistinguishable by the consensus method.

**FIG. 4:**
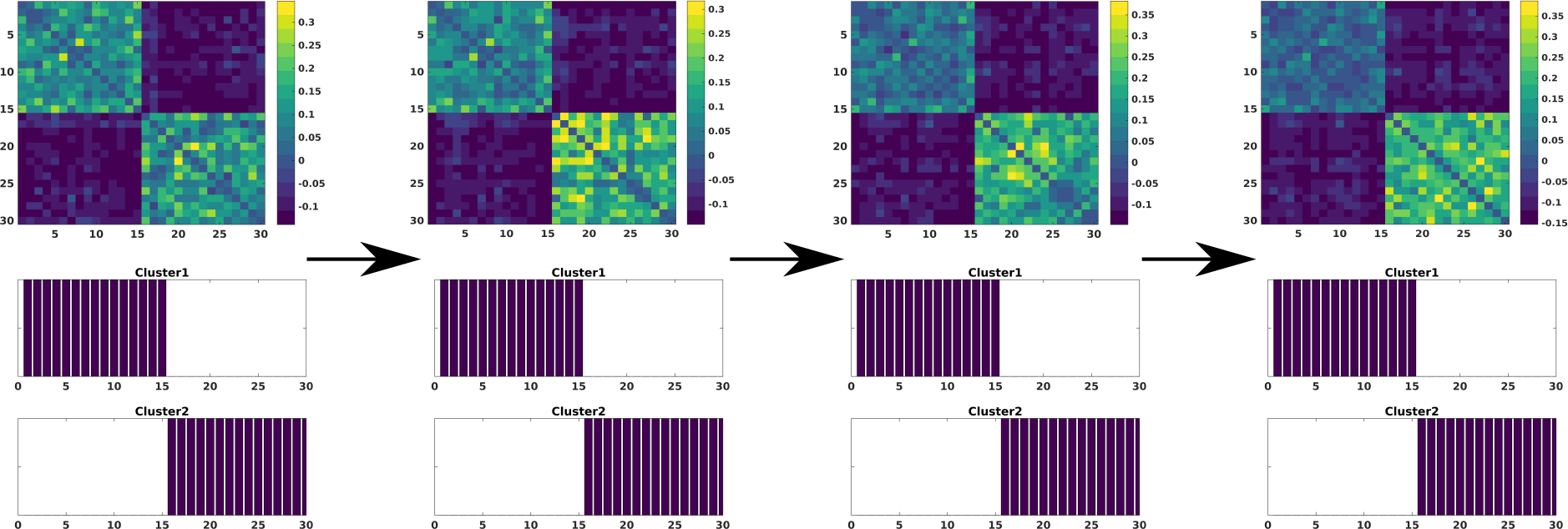
Comparison of the evolution of the modularity matrix *B* and group reconstruction when changing the subjects affected by large amplitude noise

### B. LA5C dataset

The application of the consensus clustering method to the brain matrices in the four groups yields the consensus matrices depicted in figure 5, which have been ordered according to the membership configuration found by the community Louvain routine. For the healthy group, this algorithm detects three communities of 55, 54 and 8 respectively. For ADHD group, it finds 2 clusters of 21 and 15 subjects. For BD group, it detects 3 clusters of 17, 21, 8 subjects and for SCH group 3 of 14, 25 and 9. In all cases, communities with fewer subjects than 5 have considered as outliers.

Likewise, the consensus clustering algorithm allows to extract more homogeneous subgroups of subjects, as it can be noticed from figure 5, where the mean intra-group distance distribution produced by the collection of distance matrix for each pattern connectivity is exhibited. We stress that the inter-subjects distance is highly significantly smaller for the clusters w.r.t the whole groups, as estimated by means of a Wilcoxon rank-sum test.

**FIG. 5:**
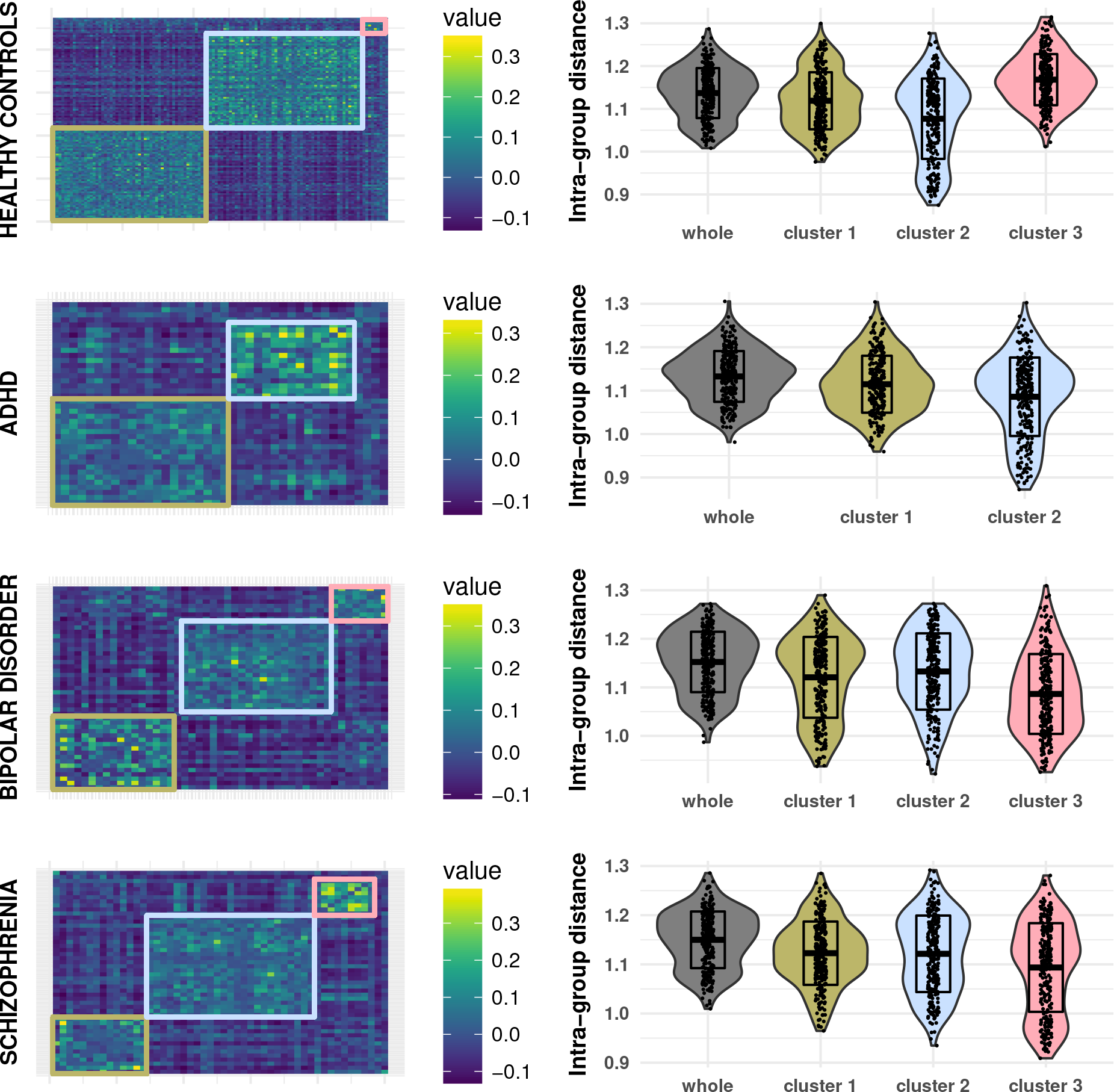
Consensus matrices and the reduction of the mean intra-group distance per node after applying the consensus clustering method

Furthermore, having more homogeneous clusters is translated in a variability decrease which leads in some cases to a gain in separability when clusters are introduced in between group comparison studies. In figure 6, we show this by means of the empirical cumulative distribution function for the uncorrected p-values from each possible comparison between whole and cluster comparison within a healthy versus pathologic group setup. As we can see, the comparisons when no clustering has been performed (represented as thick lines) begin to accumulate larger p-values in contrast to the case when some of the clusters are involved. Concerning partition of the healthy group, we can see that a procedure selecting and recruiting of subjects with similar characteristics of those subjects within the cluster 2 in our healthy control sample would optimise the node association with the pathology. In addition, ADHD, which lacks of significant results when being involved in group-comparisons as a whole, develops significance gain improvement when clusterised using the consensus algorithm. In contrast, for both BD and SCH clusters such an improvement only takes place to some of their clusters and as a consequence, the consensus algorithm can also help filter out”noisy” subjects which have less in common with the pathology in study.

**FIG. 6:**
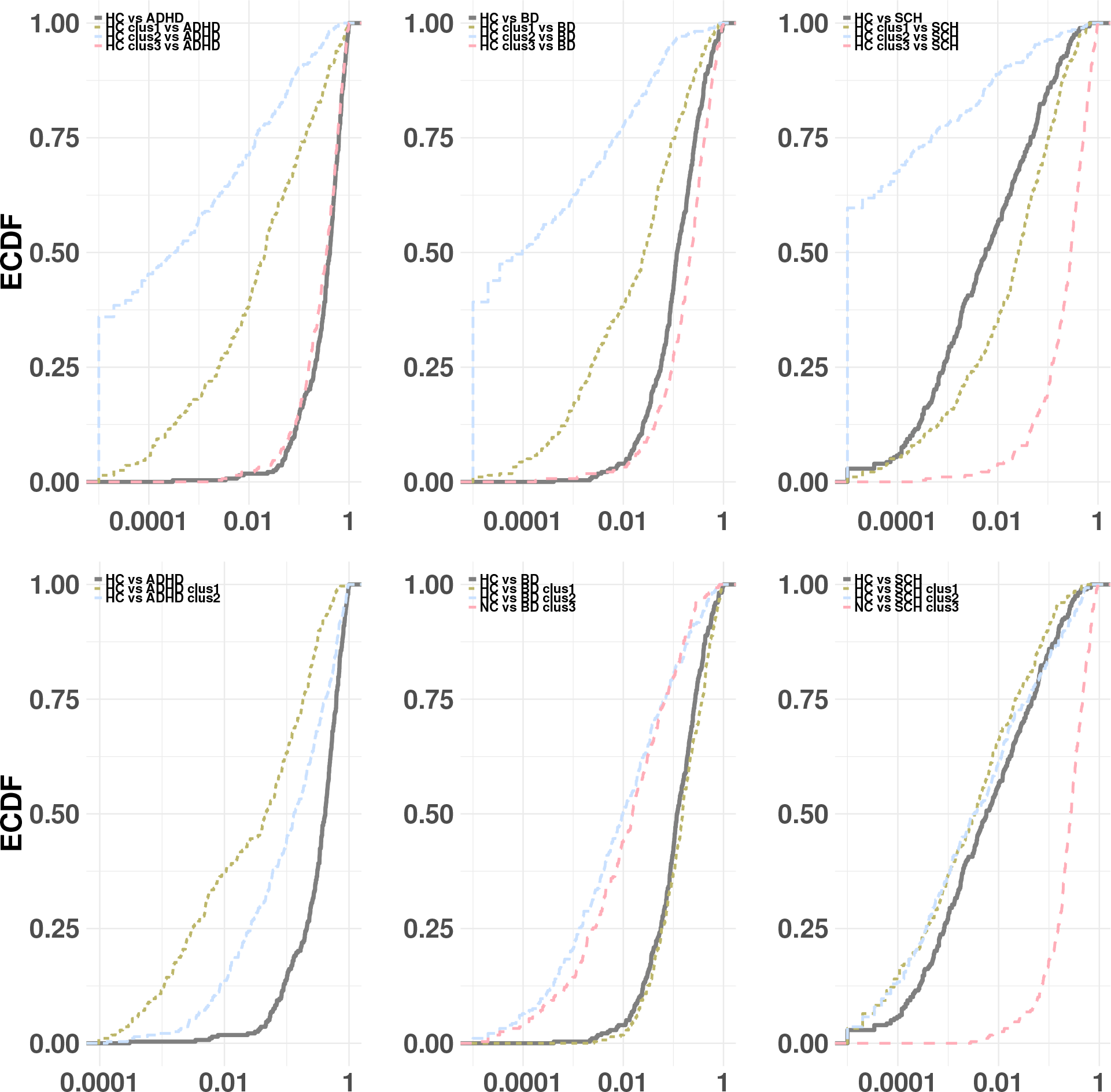
Empirical cumulative function for the uncorrected p-values in all possible comparisons involving on partitioned group

**FIG. 7:**
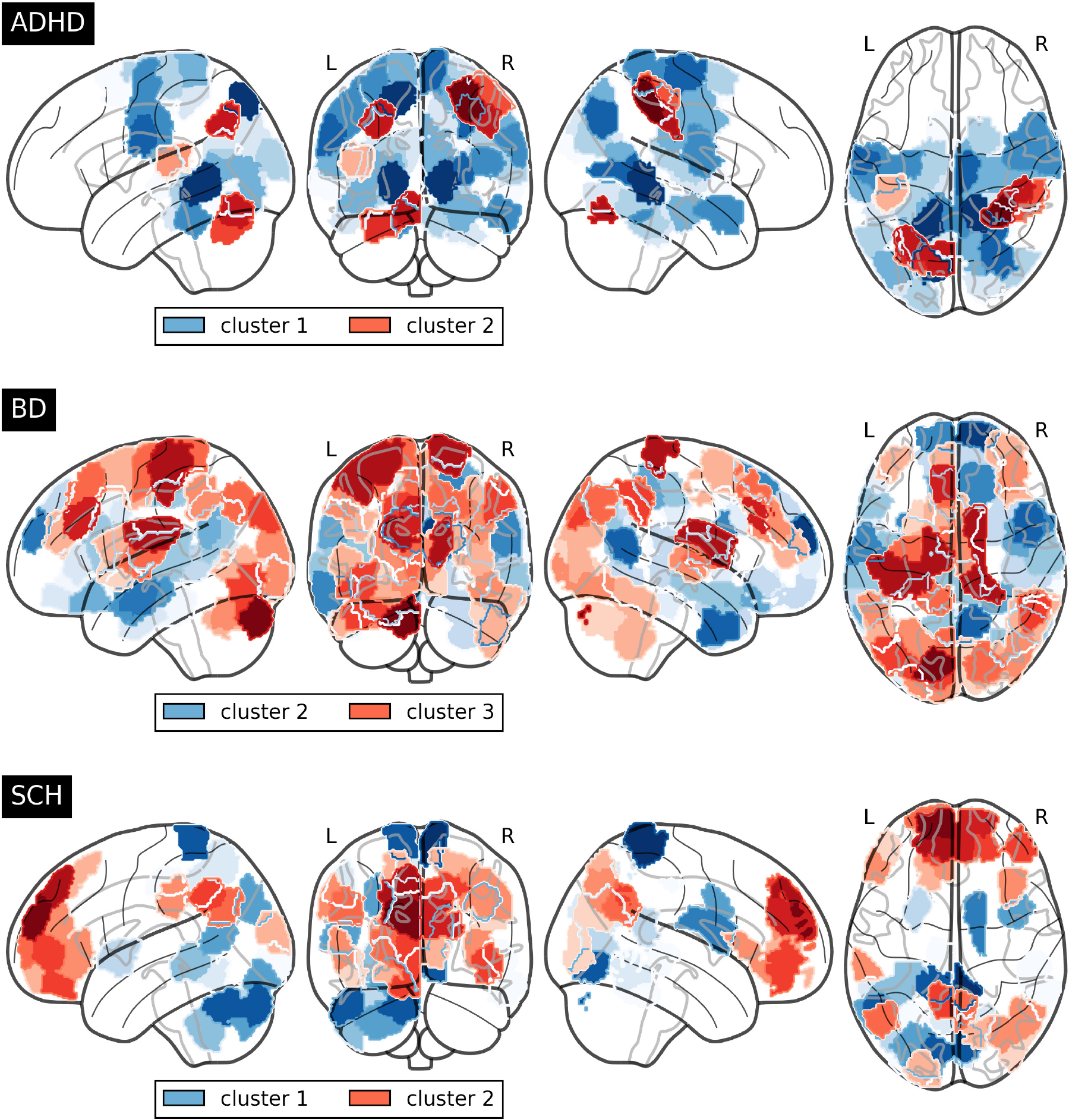
Unique node for clusters in each disorder

On the other hand, application of consensus clustering to brain connectivity matrices of pathologies can also help unmask substructures whose underlying properties are unique and that might be worth studying. For example, concerning the comparisons with healthy subjects of the ADHD group after been partitioned, we obtain for cluster 1 59 significant nodes that are not present neither in cluster 2 nor when this group is taken as a whole; we will denote these nodes as”unique nodes” for cluster 1, meaning that only the comparison with cluster 1 leads to point out their altered pattern w.r.t. pathological conditions. Likewise, 7 nodes are unique for cluster 2 of the same disorder group. Regarding comparison with the BD group, there are 61 unique nodes which are recognized as significant for cluster 1 and 46 for Cluster 2. Finally, 113 nodes arise as being significantly associated with the whole group of schizophrenia, whereas substructure inspection increases this number to 128 and 132 for cluster 1 and 2, exhibiting both 27 different unique nodes. Both cluster 1 of BD group and cluster 3 of SCH keep subjects with associated with both pathologies and therefore no association is found whatsoever. Regions affected by unique nodes in each pathology is represented in figure 7.

Higher significant regions for cluster 1 of ADHD lie in cortical regions from the Lateral Occipital Cortex, superior division, the Cingulate Gyrus, posterior division and Lingual Gyrus and involve functions from the visual and dorsal attention system. Cluster 2 have also incidence from parts of the lingual gyrus in addition to postcentral Gyrus, the superior parietal Lobule and the supramarginal Gyrus, anterior division-all with roles attached at the dorsal attention system-and a small portion of the cerbellum.

On the other hand, regarding the BD group, the default mode system fundamentally emerges in cluster 2 with cortical regions in the Frontal pole, temporal pole, the Inferior Temporal Gyrus, anterior division and the Precuneus. Cluster 3 includes the postcentral gyrus and the left thalamus, which take part in sensorimotor functions, and parts of the cerebellum and the Occipital Fusiform Gyrus.

Finally, SCH community representation obtained through the consensus algorithm exhibit a well differentiated structure, with cluster 1 focusing mainly on the back parts of the brain with the Postcentral Gyrus, Precuneous Cortex and Lingual Gyrus and play roles from the somatosensory and visual systems, and cluster 2 involving basically the prefrontal cortex and some parts of the Superior frontal Gyrus whose functions can be assigned to the default mode network.

**FIG. 8:**
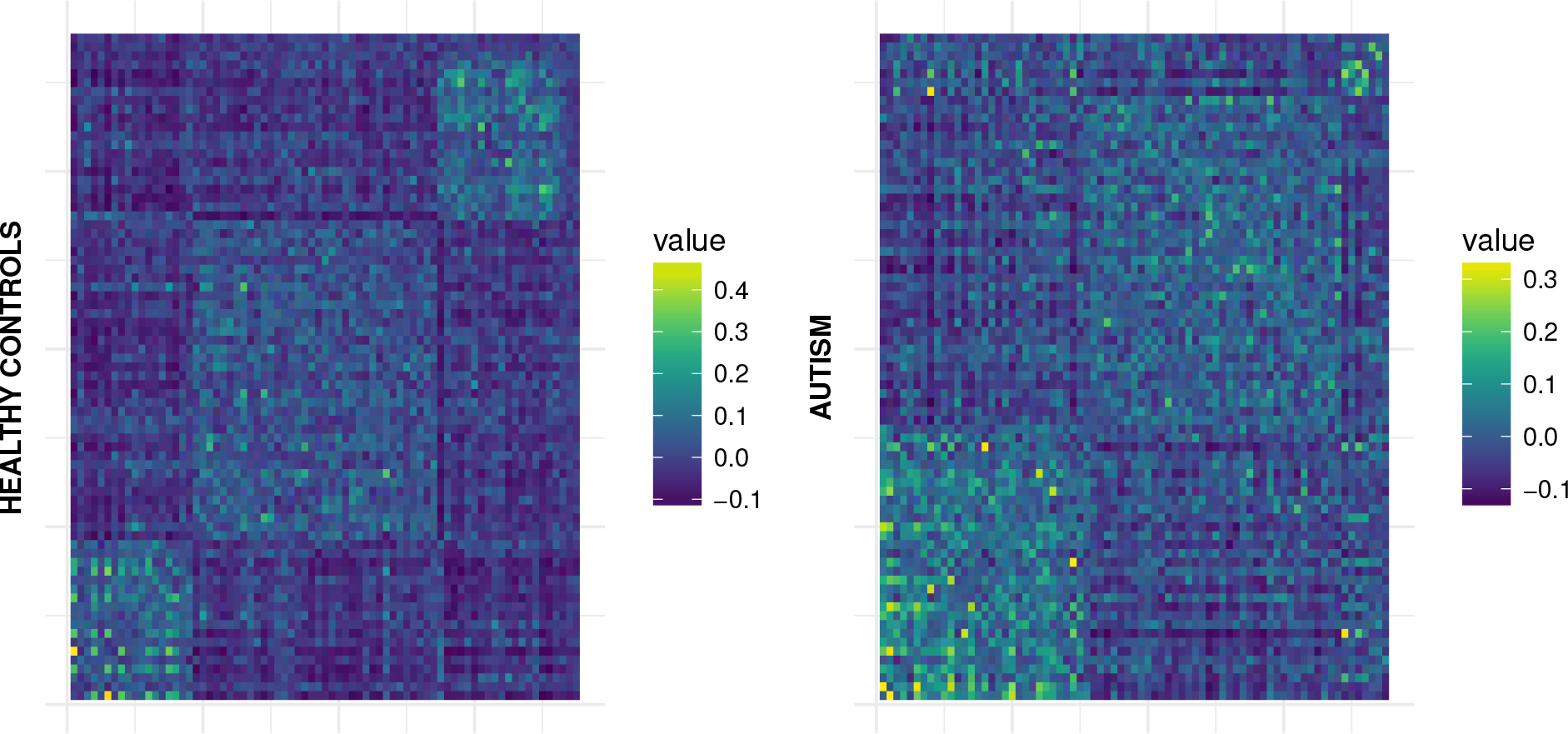
modularity matrix *B* ordered after label structure provided by community detection for healthy (left) and autistic (right) group

## IV. ABIDE DATASET

Application of the consensus clustering algorithm and subsequent community detection algorithm to both groups leads to the ordered community structure provided by the consensus matrices in figure 8. For healthy group, the modularity matrix *B* yields 3 modules of 18, 36 and 19 subjects respectively. In contrast, division of autism group consists of two modules of 30 and 38 patients respectively. Moreover, a Wilcoxon Rank sum test performed on ASD community partition separates subjects statistically by age (*p* = 0.0028) and framewise displacement (*p* < 0.0001) (see fig 9).

On the other hand, we can see in figure 10 that the statistical significance is overall stronger when a partitioned case is involved. In particular, it is notable how cluster 3 of healthy group shows all the benefits of applying the consensus clustering method since it gathers the healthy subjects with the highest connectome difference in comparison with autistic subjects. Moreover, this pronounced case corresponds to the scenario where the decrease of intra-group distance (cluster is more homogeneous) also follows an increase of the inter-group distance (left panel of figure 11). This also happens less notoriously for cluster 1 of autism group. In the rest of the cases, even though clusters become more compact, such a behaviour does not take place, given that, since we are clustering each group separately, maximum separation between classes does not need to be guaranteed. Nevertheless, the gain in statistical significance for some of the case might allow to unmask regions not observed significantly different at first and therefore be further exploited by a more thorough exploratory analysis.

**FIG. 9:**
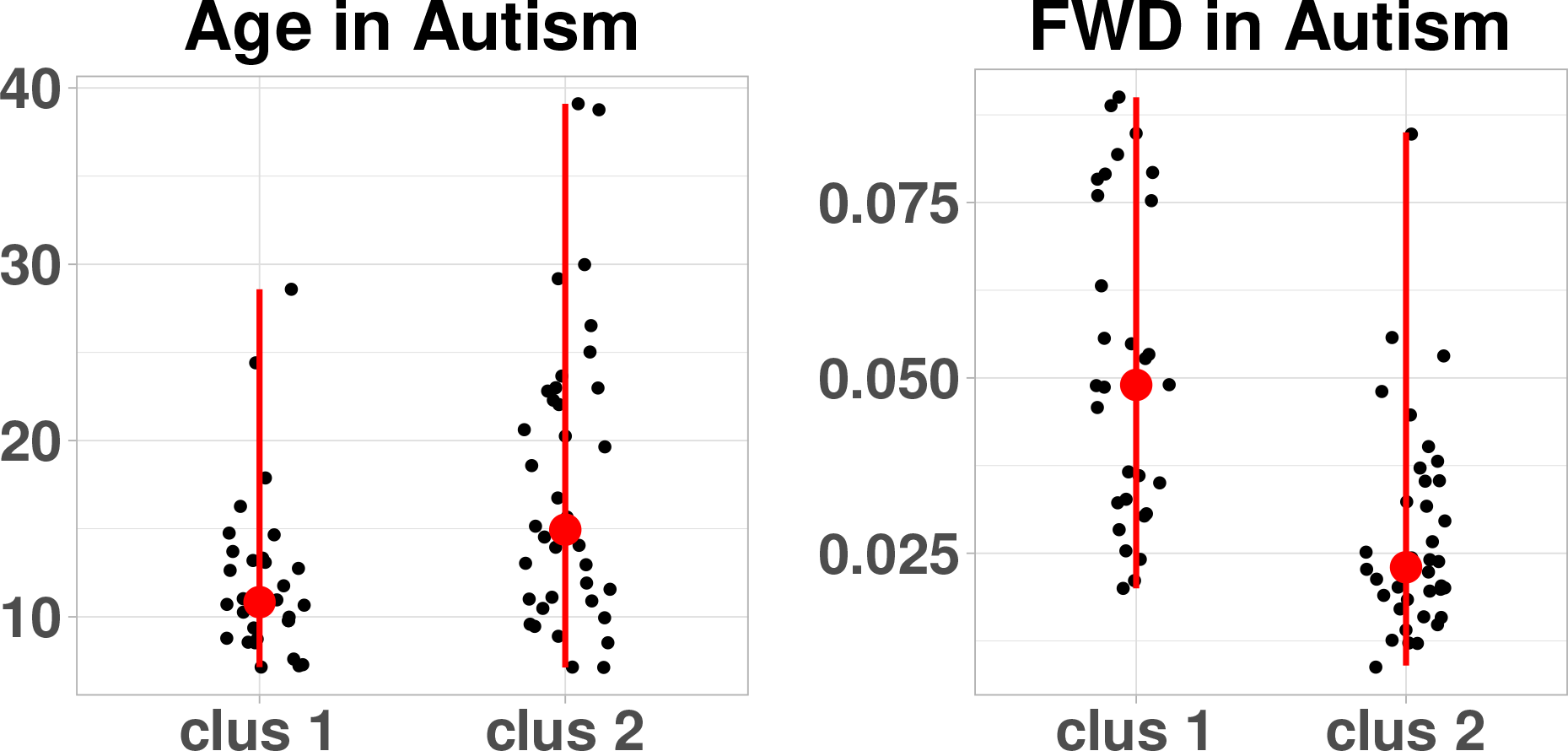
Age and FWD distribution for ASD clusters

When analysed together both groups each as a whole, MDMR yields only 2 significant regions (region 57 and 83) as explained by group-label variation between subjects, whereas this number increases to 13 and 70 for age and FWR respectively, showing an important effect of the observed variance based on these last variables. However, applying first consensus clustering to the healthy group produces 3 clusters such that, when comparing with the ASD group, one cluster yields 52 significant nodes, being 48 not in any other possible comparison. Likewise, both clusters obtained by partitioning ASD group lead to 16 and 9 significant nodes when compared with healthy group and most importantly, 14 and 6 of them respectively only observable in each cluster. Therefore, not only do we gain separability by means of an increase of significant nodes, but also our approach allows us to unmask unique regions not visible whatsoever when using standard whole groups comparisons.

From a clinical point of view, the emergence of unique nodes can be important and lie in brain regions whose implications on the development of the pathology should be further examined. In our case, concentrating on the ASD case, cluster 1 has 14 regions not presented neither in cluster 2 versus HC, nor in whole group comparison. These regions are specially focalised on the right brain hemisphere with the highest overall significance located at the pars triangularis of the inferior frontal gyrus (*p*^fdr^ = 0.0004). Areas in the same hemisphere involving the frontal pole (*p*^fdr^ = 0.0018) and the precuneus (*p*^fdr^ = 0.0036) and regions from the the postcentral (*p*^fdr^ = 0.0014) and paracentral (*p*^fdr^ = 0.0019) in the opposite hemisphere exhibit also a great significance. On the other hand, the 6 unique significant nodes of cluster 2 belong to areas in both hemispheres involving the Banks of the Superior Temporal Sulcus (*p*^fdr^ = 0.0021 in the left hemisphere and *p*^fdr^ = 0.0078 in the right) and the superior temporal lobe (*p*^fdr^ = 0.0085 and p^fdr^ = 0.0075 respectively); and regions in the inferior temporal lobe (*p*^fdr^ = 0.0069) and the supramarginal (*p*^fdr^ = 0.0079) in the right hemisphere.

**FIG. 10:**
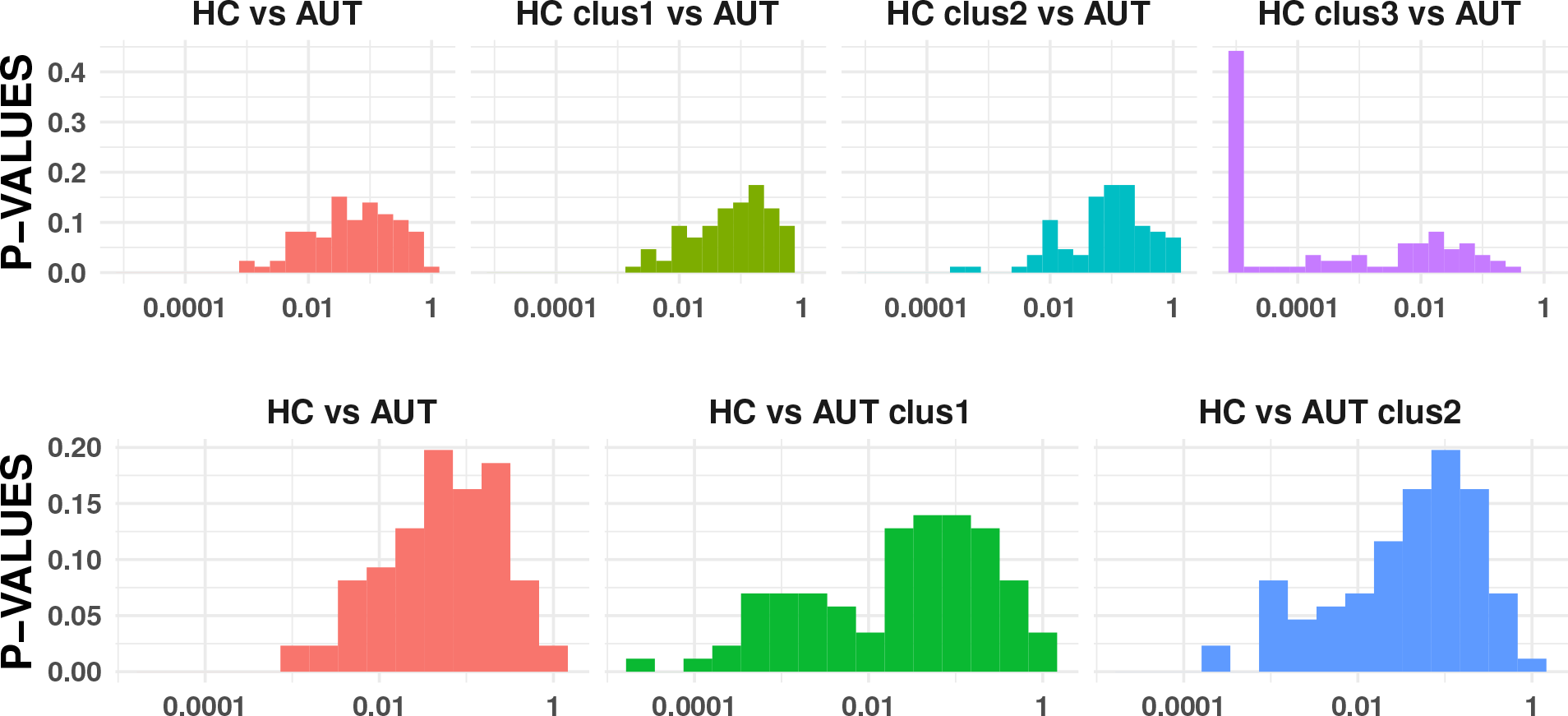
Uncorrected p-values distribution for all possible comparisons involving one partitioned ABIDE group from the application of the consensus clustering

## V. DISCUSSION

The incorporation of new techniques for increasing separability in neuroimaging studies is important for the obtainment of new findings. Our clustering method deals naturally with brain connectivity matrices, providing underlying more compact sub-classes beyond the usual group labels. In addition, the application of the consensus clustering to different regions of the brain allows us to capture the differences in groups in comparison to using the whole connectivity matrix. To our best knowledge, this has not been considered yet at least in the neuroimaging field. On the other hand, consensus matrix gathers all the information from different resolutions and nodes and encodes this in the consensus matrix. As a consequence, arbitrariness brought about by a priori choice of the number of clusters disappears. Moreover, clusters found by our method are fairly robust with respect to moderate changes in amplitude noise. However, given nowadays the strong protocols in experiment design and the successive denoising steps in the preprocessing pipeline, one would not expect to find such a noise contaminating the real data.

**FIG. 11:**
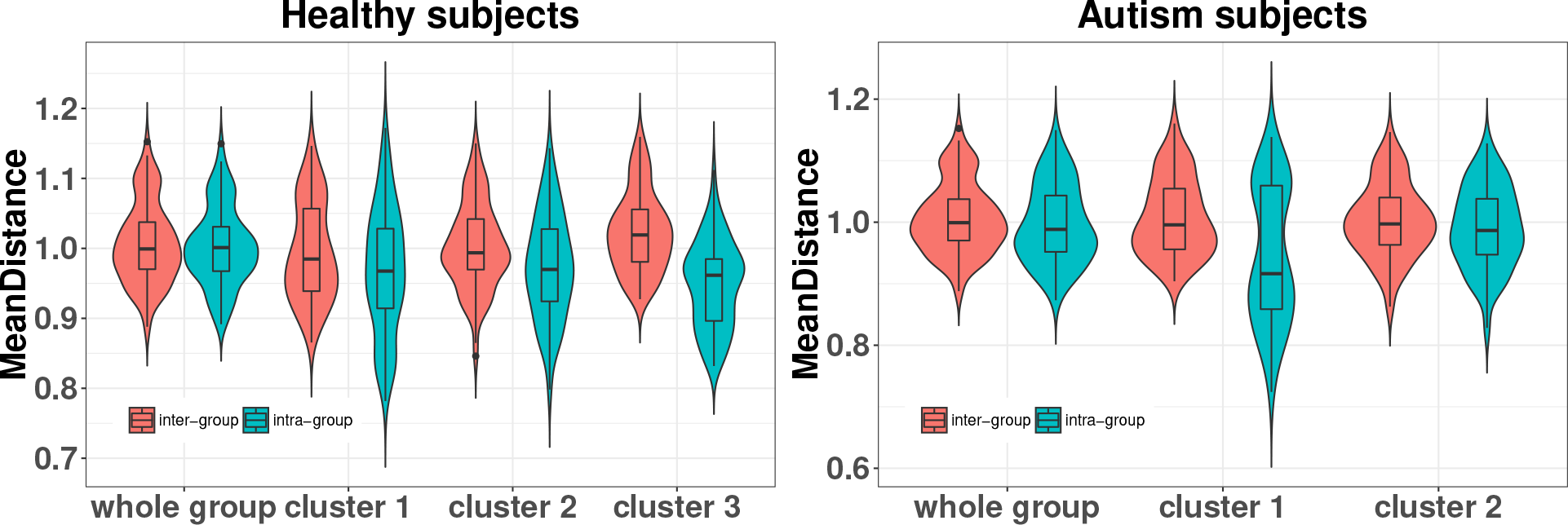
mean intra and inter-group distance distribution by the collection of distance matrices per each node

In addition to variability reduction that allows to elucidate significant pattern connectivity regions in group comparisons, each cluster found also provides unique information. In particular, we have shown this for two different datasets which include four different mental disorders: ADHD, BD, SCH and ASD.

Most of the studies on finding brain alterations in ADHD subjects have been focused on children, since the symptoms related to this disorder such as inattention, and/or hyperactivity/impulsivity need to be treated since the beginning of their appearance. In our case, however, our cohort consists of adult subjects and we have found two different clusters carrying both different and unique information. The first cluster comprises a vast anterior brain area, with Lateral Occipital Cortex and the lingual gyrus and are mainly responsible both for visual attention. This has already been reported using structural data [20]. The second one covers a small portion of the brain developing functions from the dorsal attention and the cerebellum, maybe capturing the cognitive functions of this last structure in attention tasks [21].

Amongst our findings, our clustering method clearly separates the Default Mode Network as one of the relevant system embedded in one of the subgroups in Bipolar Disorder and Schizophrenia. Such a fundamental network has already been reported many times and it is known to play a crucial role in both mental disorders [22, 23]. In our specific clusters, regions with a disrupted connectivity pattern concentrates on the frontal cortex [24, 25], with also some areas in the temporal lobe with prominence in bipolar disorder subjects, which may account for differences when compared with schizophrenia [26–28].

Interestingly, our method also allows us to provide areas involved in different functional systems, such as the cerebellum [29–31], the appearance of delusion and hallucinations conducted by disfunction of sensory system located in the postcentral gyrus [32] and lack of empathy triggered by an hyperactivation of the bilateral lingual gyrus in schizophrenic subjects [33].

On the other hand, our results regarding ASD subject show two clear region domains. The first one with a clear right asymmetry, with the participation of areas around the Pars opercularis and Pars triangularis in the Inferior frontal gyrus, and whose individuals usually exhibit lack of emotions [34-36]. Likewise, this domain also includes sensory-motor areas such as the postcentral and paracentral gyrus and precuneus and confirm the results observed in a larger ABIDE cohort of subjects when comparing ASD with typically developing controls [37]. Second, some parts of the temporal lobe which, in addition to regions in the orbito-frontal cortex (OFC) and amygdala define the”social brain”. In this case, autistic subjects specialise for verbally labelling complex visual stimuli and processing faces and eyes, which compensate amygdala abnormality [38].

It is worth to stress that the connectivity patterns among brain regions are influenced by respiration, movement, cardiac phase and other physiological variables, and what we observe and can infer here (and in most of fMRI studies) is just a higher level view in which all these actors are conflated.

## VI. CONCLUSIONS

In this work we have proposed the use of the consensus clustering approach developed in [7] for exploratory analysis, in order to cope with the hetereogeneity of subjects. Extracting the natural classes present in data and subsequently performing the supervised analysis by MDMR, between the subgroups found by consensus clustering, allows to identify variables whose pattern is altered in group comparisons, and which are not highlighted when the groups are used as a whole. As a result, the proposed approach leads to an increase in statistical power. We present application of the proposed method to fMRI public data, looking for brain regions whose connectivity pattern is altered in the group comparisons which are significantly altered due to the pathology.

